# A *space-jump* derivation for non-local models of cell-cell adhesion and non-local chemotaxis

**DOI:** 10.1101/093617

**Authors:** Andreas Buttenschön, Thomas Hillen, Alf Gerisch, Kevin J. Painter

## Abstract

Cellular adhesion provides one of the fundamental forms of biological interaction between cells and their surroundings, yet the continuum modelling of cellular adhesion has remained mathematically challenging. In 2006, Armstrong *et al.* proposed a mathematical model in the form of an integro-partial differential equation. Although successful in applications, a derivation from an underlying stochastic random walk has remained elusive. In this work we develop a framework by which non-local models can be derived from a space-jump process. We show how the notions of motility and a cell polarization vector can be naturally included. With this derivation we are able to include microscopic biological properties into the model. We show that particular choices yield the original Armstrong model, while others lead to more general models, including a doubly non-local adhesion model and non-local chemotaxis models. Finally, we use random walk simulations to confirm that the corresponding continuum model represents the mean field behaviour of the stochastic random walk.

## 1 Introduction

The building blocks of multicelluluar organisms are cells, vessels and protein fibres. These tissue constituents display complex biochemical and physical interactions with cell adhesion and chemotaxis being two such examples. Adhesion is facilitated by transmembrane adhesive complexes, while chemotaxis requires receptors on the surface of the cell membrane. Both chemotaxis and cellular adhesion are instrumental in embryogenesis, cell-sorting and wound healing, along with homoeostasis in multicellular organisms. Misregulation of chemotaxis and cellular adhesion may lead to pathological conditions such as cancer and other degenerative diseases (Friedl and Alexander, 2011; Roussos et al., 2011): for instance, reduced cellular adhesion in cancer cells can lead to invasion and metastasis (Friedl and Alexander, 2011). Both chemotaxis and cellular adhesion have been the focus of intense mathematical modeling efforts, both at the level of agent-based and continuum models, with the latter typically based on local or non-local partial differential equations (Armstrong et al., 2006; Domschke et al., 2014; Hillen and Painter, 2009; Horstmann, 2003; Painter, 2009). Local models, typically based on the reaction-advection-diffusion framework, have a solid foundation in biased random walks. Non-local models appear to be better suited to describe certain aspects of adhesion, along with non-local chemotaxis, yet as far as we are aware there is no convincing microscopic random walk process that leads to these non-local models. In fact Gerisch and Painter (2010) have written on page 328 that “A highly desirable objective is to develop continuous models for cellular adhesion as the appropriate limit from an underlying individual model for cell movement”. In this paper we propose to fill this gap and introduce a spatial stochastic random walk that leads, in an appropriate limit, to the non-local adhesion and chemotaxis models. This approach provides a better understanding of the underlying modelling assumptions and allows to modify the continuous model as needed.

### 1.1 Biological Background

This section reviews pertinent biological background on cell adhesion, required for the subsequent derivation. Cellular adhesions are facilitated by cell adhesion molecules, which are proteins present on the cell surface (Leckband, 2010). In layman’ s terms, they act to stick cells to each other and their surroundings. Through these connections, cells can sense mechanical cues and exert mechanical forces on their environment. It is of note that cellular adhesion is important in both adherent (static) cells and in motile cells (Leckband, 2010). Due to their importance adhesion molecules are ubiquitous in biological organisms and, accordingly, it is not surprising that numerous adhesion molecules are known with integrins and cadherins forming two prominent classes. Integrins are important in both the static and dynamic case (Leckband, 2010) and are commonly associated with contacts between cells and the extracellular matrix (ECM) (Desgrosellier and Cheresh, 2010). Cadherins, on the other hand, are more commonly associated with forming stable cell-cell contacts (Leckband, 2010). The density and types of presented adhesion molecules determine the mechanical interaction strength between a cell and its neighbours or the environment. While cell adhesion is important during both homoeostasis and during cell migration, we will treat the homeostatic case as a special case of migration, in which cells are in a mechanical equilibrium or at a so called steady state.

The motility of cells is fundamental during many biological functions, including embryogenesis and wound healing (Davies, 2013). In this dynamic setting, cellular adhesion plays an important role in guiding migrating cells. Cells have developed many mechanisms of translocation, with at least four different kinds of membrane protrusions distinguished: lamellipodia, filopodia, blebs and invadopodia (Ridley, 2011). How the directionality of these protrusions is determined varies greatly between cell types, but a unifying feature of migrating cells is the formation of a spatially asymmetric morphology, allowing a clear distinction between front and back (Lauffenburger and Horwitz, 1996; Ridley et al., 2003). This is the so-called process of cell *polarization* and can result from a wide variety of intrinsic and extrinsic cues, including chemical or mechanical stimuli (Danuser et al., 2013; Geiger et al., 2009; Ridley et al., 2003; Théry et al., 2006; Weiner et al., 2000). Once polarized, membrane protrusions are extended primarily at the cell front (Lauffenburger and Horwitz, 1996).

### 1.2 Mathematical Background

Many partial differential equation models for the dynamics of cellular populations are motivated by the conservation of mass equation. Suppose that the population density is given by *u*(*x, t*) and its flux by *J* (*x, t*). The conservation equation is then given by,

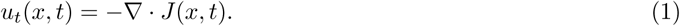

Different biological phenomena can be described by an appropriate choice for the flux. In many common models the flux is divided into multiple additive parts: for example, a part due to random motion denoted *J*_*d*_, and a part due to adhesion denoted *J*_*a*_, that is

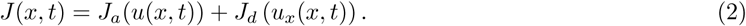

In the following discussion we focus on different choices for *J*_*a*_ and assume Fick’ s law for the diffusive flux, i.e., *J*_*d*_ = *-Du*_*x*_(*x, t*). Armstrong et al. (2006) suggested the following flux term to model movement through the formation and breaking of cell-cell adhesion between cells. The flux *J*_*a*_ is assumed to be directly proportional to the created force and the population density, and inversely proportional to the cell size, such that

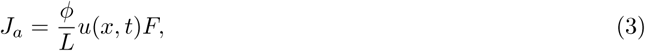

where ϕ is a constant of proportionality, *F* the total adhesion force and *L* the typical cell size. This follows from Stokes’ law, which gives the frictional force of a spherical body moving at low Reynolds numbers. The total adhesion force is the result of the forces generated within the sensing radius, i.e.,

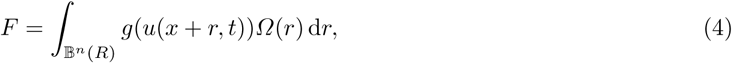

where *g*(*u*(*x, t*)) describes the nature of the force and its dependence on the cell density and *Ω*(*r*) describes the force’ s direction and dependence on the distance *r*. Substituting the fluxes into the conservation equation we obtain the following integro-partial differential equation:

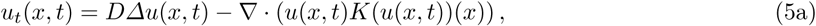

where

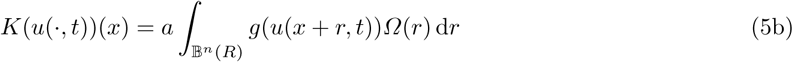

and *a* = ^*ϕ*^*/*_*L*_. It is noted that this model was the first continuum model that successfully replicated the patterns observed in Steinberg’ s classic adhesion-driven cell-sorting experiments (Armstrong et al., 2006).

In this paper we will focus on the modelling of the non-local adhesion process, for this reason we assume in equation (5) that the diffusion coefficient is constant. For the modelling of spatial dependent diffusion coefficients we refer the reader to (Stevens and Othmer, 1997) and (Calvo et al., 2015).

The adhesion model (5) has since been extended to include both cell-cell and cell-ECM adhesion (Gerisch and Chaplain, 2008), with this extended version used to study the invasion of cancer cells into the ECM (Andasari et al., 2011; Chaplain et al., 2011; Domschke et al., 2014; Gerisch and Chaplain, 2008; Painter et al., 2010; Sherratt et al., 2008). The adhesion model was also used in a model of neural tube development in vertebrates (Armstrong et al., 2009). In addition to the various biological applications, the mathematical properties of the adhesion model (5) have been under intense investigation: conditions for global existence were developed in (Chaplain et al., 2011; Dyson et al., 2010; Sherratt et al., 2008; Winkler et al., 2016); travelling wave solutions were found for small parameters *α* in equation (5b) (Ou and Zhang, 2013); an efficient and robust numerical scheme for the non-local adhesion model has been developed in (Gerisch, 2010).

While successful in application it is not yet clear how the microscopic properties of cells, such as cell size, cell protrusions, nature of the adhesion forces or the distribution of adhesion molecules enter the adhesion model (5). A derivation of the non-local adhesion model (5) from an underlying random walk promises to answer these questions. An early attempt was undertaken by Turner et al. (2004), who studied the continuum limit of a Cellular Potts model that included cell-cell adhesion, yet the resulting continuum model was too complicated for significant analysis. Johnston et al. (2012) studied the continuum limit they obtained from a stochastic exclusion movement model; under certain parameter regimes, however, the continuum model permits negative diffusion and is unable to replicate the pattern formation observed during cell-sorting. Recently, Middleton et al. (2014) showed that an integro partial differential equation (iPDE) model similar to equation (5a) can be obtained from the mean field approximation of Langevin equations. There it is shown that this iPDE limit is only effective for weak cell-cell adhesion.

Many commonly used partial differential equation models, such as the chemotaxis equation, have been derived from a space-jump process (Hillen and Painter, 2009; Othmer et al., 1988). This motivates us to derive the adhesion model (5) from an underlying stochastic random walk model based on cell-cell mechanics. The challenge is maintaining the finite sensing radius while taking the formal limit of the space-jump model. Through this we offer not only insight into the complicated assumptions on which Armstrong’ s model (5) is based, but also extend it to a more general form. We see that Armstrong’ s model implicitly assumes (i) point-like cells, (ii) mass action for the adhesion molecule kinetics and (iii) no volume exclusion effects. Our approach allows us to relax these assumptions and extend the model to include large cells, volume exclusion and more complicated adhesion molecule kinetics. In the process we find that our derivation is general, and hence applicable to other non-local models such as the non-local chemotaxis model in Section 4.

### 1.3 Layout of the paper

The key to a mathematical derivation of a continuum adhesion model lies in the definition of cell polarization and in Section 2 the biological notion of a polarization vector is integrated into the framework of a stochastic random walk. This preparatory work will allow us in Section 3 to derive the non-local adhesion model from an underlying stochastic random walk and in Section 4 to derive the non-local chemotaxis model. In Section 5 the proposed stochastic random walk is compared to numerical simulations of the integro-partial differential equation model of adhesion. In the final Section 6 our results will be discussed and an outlook given on open questions and future work.

## 2 A population model informed by the polarization vector

For the following discussion we let the domain be given by *Ω*. Here we restrict the domain to *Ω* = ℝ ^*n*^ in order to focus on modelling cell-cell interactions in the absence of boundary effects. Suppose that a cellular population with population density function *u*(*x, t*) is present in *Ω*. To define precisely the meaning of this population, we make a couple of assumptions. Cells send out membrane protrusions to sample their environment and we assume that these membrane protrusions are much more frequent than translocations of the cell body; the cell body includes the cell nucleus and most of the cell’ s mass. In modelling cell migration we are most interested in the translocation of the cell body and not the frequent, but temporary, shifts due to cell membrane protrusions. For this reason, we define the population density function *u*(*x, t*) as follows,

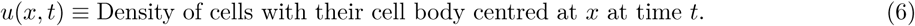

This precise definition of the cell population will become important later in Section 3.

We model the evolution of the cell population u(x, t) using a space-jump process, using stochastically independent jumpers. In other words, we use a continuous time random walk for which we assume the independence of the waiting time distribution and spatial redistribution. Here the waiting time distribution is taken to be the exponential distribution, with a constant mean waiting time. A possible extension in which the mean waiting time is a function of cell density is briefly motivated in Section 6. Let *T* (*x, y*) denote the transition rate for a jump from *y* ∈ *Ω* to *x* ∈ *Ω*. The evolution of the population density *u* (*·, t*) due to these particle jumps is given by the Master equation (Othmer et al., 1988),

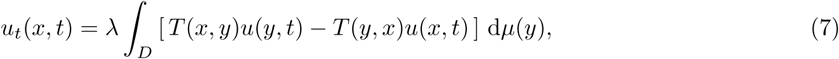

where (*D, µ*(*y*)) is a measure space with *D ⊂ Ω*, and 1*/λ* is the mean waiting time. Note that, *T* (*x, y*) depends on the population density *u*(*·, t*), however we do not explicitly track this to keep the notation managable and within convention see for instance (Hillen and Painter, 2009). For more details on the derivation of the master equation see (Hughes, 1995; Van Kampen, 2011).

We make two assumptions about the movement of the cells:

### Modelling Assumption 1

We assume that in the absence of spatial or temporal heterogeneity the movement of individual cells can be described by Brownian motion. It has been shown that this is a reasonable assumption for many cell types (Beysens et al., 2000; Mombach et al., 1995; Schienbein et al., 1994)

### Modelling Assumption 2

The cells’ polarization may be influenced by spatial or temporal heterogeneity. We denote the polarization vector by **p**(x)

In this section we will show how these two assumptions can be naturally included within the space-jump framework. We will further discuss the formal limit of equation (7) which will lead to macroscopic models describing the spatial-temporal evolution of *u*(*x, t*). Figure 1 gives an overview of how the polarization vector and the diffusion scaling are used to obtain the final population model.

**Fig. 1:**
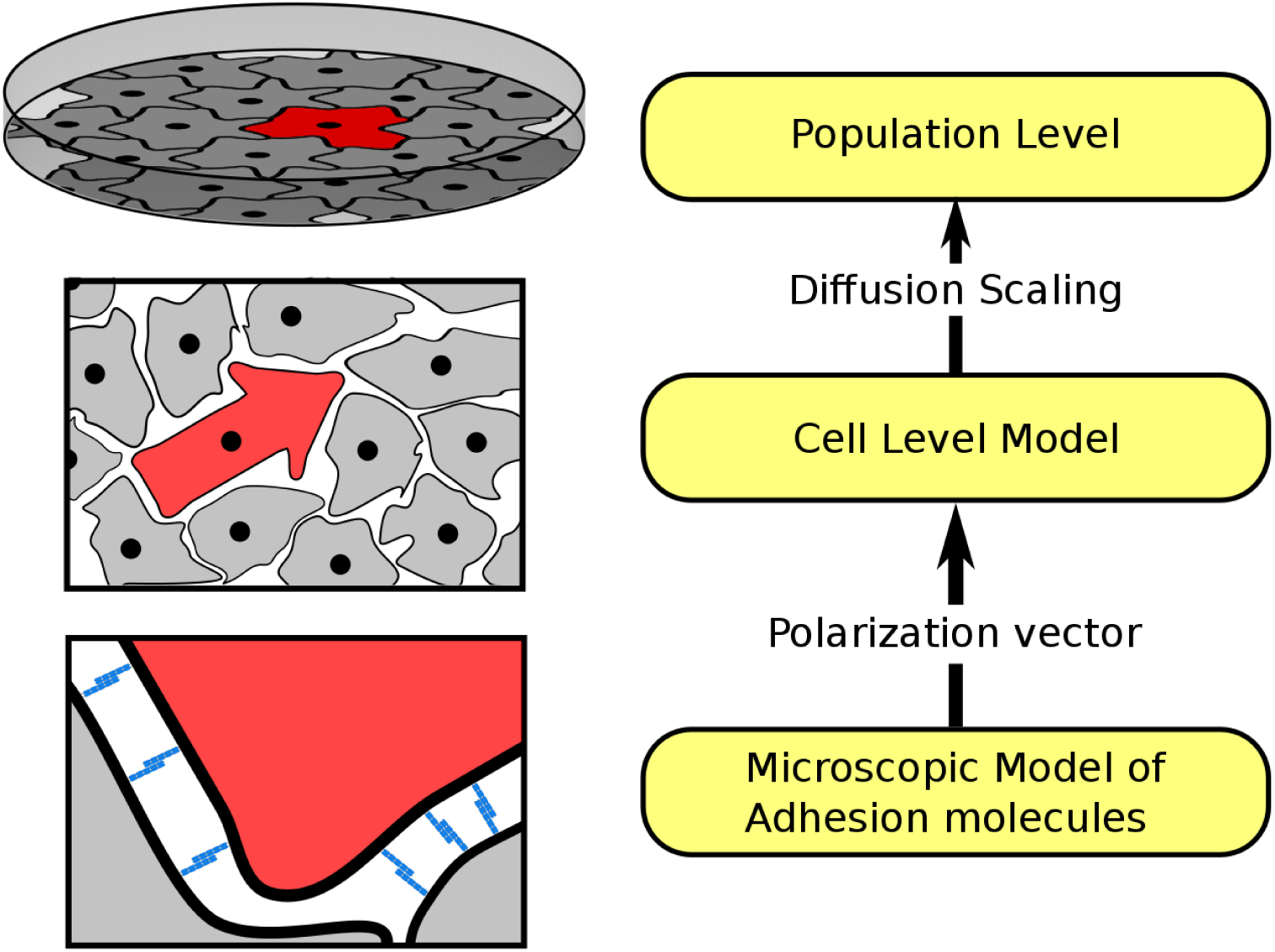
Schematic of the bottom-up approach of modelling cell movement due to cell-cell adhesion. Our final goal is to derive an equation describing the evolution of the cell population density. This is called the *population level*. The population level model is derived from a *cell level model*. Finally, the cell level model is informed by the *polarization vector*. The cell depicted in red is given an arrow shape to indicate the polarization vector **p**(*x*). The polarization vector is determined by the formation and breaking of adhesion bonds between the cells. The adhesion bonds are depicted as blue sticks. The polarization vector connects the microscopic and the mesoscopic scale (see Section 2.1). The diffusion scaling connects the mesoscopic and the population level (see Section 2.3).

#### 2.1 The polarization vector in a space-jump process

For notational convenience we associate to a jump from *y∈ Ω* to *x ∈ Ω* the heading *z*:= *x - y*. Using the heading we define *T*_*y*_(*z*) =: *T* (*y* + *z, y*) = *T* (*x, y*). Let *D*^*y*^ denote the set of all possible headings from *y*. We assume that if *z∈ D*^*y*^ then so is -*z∈ D*^*y*^. In other words, *D*^*y*^ is a symmetric set. In most cases *D*^*y*^ will be independent of *y*. We further assume that for every *y ∈ Ω*, the function *T*_*y*_ is non-negative as it represents a rate.

Given *y∈ Ω* we denote the redistribution kernel at this location by *T*_*y*_(*z*); we assume that *T*_*y*_ *∈ L*^1^(*D*^*y*^) and that ‖ *T*_*y*_‖ _1_ = 1 holds. Note that here we use the measure associated with *D*^*y*^. This measure can be the standard Lebesgue measure in ℝ ^*n*^ if *D*^*y*^ is the ball with radius *h* in ℝ ^*n*^, i.e., *D*^*y*^ = 𝔹 ^*n*^(*h*), or it could be the surface Lebesgue measure in ℝ ^*n*^ if *D*^*y*^ is the sphere with radius *h* in ℝ ^*n*^, i.e., *D*^*y*^ = 𝕊 ^*n-*1^(*h*), or it can be a discrete measure if modelling movements on a lattice and *D*^*y*^ =*{ he*_1_*, −he*_1_*, he*_2_*, …, −he*_*n*_ *}*, where *e*_1_*, e*_2_*, …, e*_*n*_ are the Cartesian unit vectors. In any case, this normalization makes *T*_*y*_ a probability density function (pdf) on *D*^*y*^.

Any function which is defined for both *z* and -*z* can be decomposed into even and odd components, which are denoted by *S*_*y*_ and *A*_*y*_ respectively. This notation is chosen in imitation of the even/odd notions introduced by Mogilner and Edelstein-Keshet (1999) in a non-local modelling study of swarming behaviour.

### Lemma 1

*Consider y ∈ Ω, given T*_*y*_ *∈ L*^1^(*D*^*y*^)*, then there exists a decomposition as*

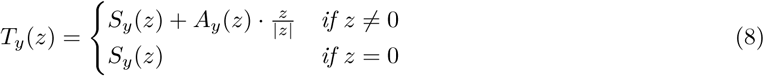

*with S*_*y*_ *∈ L*^1^(*D*^*y*^) *and A*_*y*_ *∈ (L*^1^(*D*^*y*^)) ^*n*^*. The even and odd parts are symmetric such that*

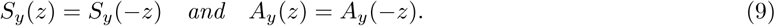

*Proof.* Given *T*_*y*_, define

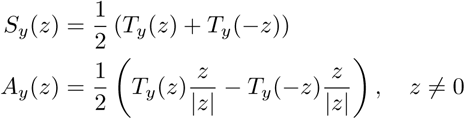

Then the above properties can be checked by direct computation.

Using this decomposition we define two properties which are analogous to Modelling Assumptions 1 and 2 above. First, we define the motility.

**Definition 1 (Motility)** We define the motility at *y ∈ Ω* as

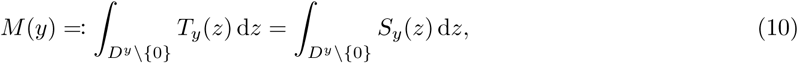

where the integration is w.r.t. the measure on *D*^*y*^.

The motility is the probability of leaving *y*. This probability is 1 if 0 *∉ D*^*y*^; it is also 1 if 0*∈ D*^*y*^ and *T*_*y*_ is a continuous pdf, and it may be smaller than 1 if 0 *∈D*^*y*^ and *T*_*y*_ is a discrete pdf. Here we find that the motility depends solely on the even component *S*_*x*_, in other words solely on modelling assumption one.

Secondly, we define the polarization vector in a *space-jump* process.

**Definition 2 (Polarization Vector)** The polarization vector at *y ∈ Ω* is defined as,

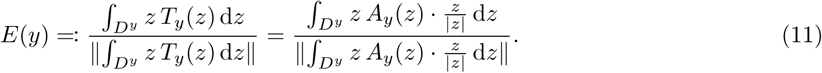

where the integration is w.r.t. measure on *D*^*y*^.

The first moment of the pdf *T*_*y*_ can be intuitively understood as the expected heading of a jump originating at *y*. This is in direct correspondence with a polarized cell which, following polarization, moves in the direction of the polarization vector. The expected heading is solely determined by *A*_*y*_, which therefore plays the role of the polarization vector **p**(*y*) in a space-jump process. This correspondence motivates us to set *A*_*y*_ = **p**(*y*) in the subsequent derivations.

#### 2.2 Derivation of macroscopic equations

For the following derivations of the population model, we require the following assumptions.

1. We consider a myopic random walk, i.e.,

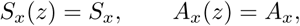

then

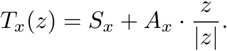 Note that *T*_*x*_(*z*) still depends on *z* and *x* and the scalar *S*_*x*_ and vector *A*_*x*_ depend on *x*.
2. We consider small jumps of length *h*, and write

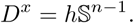 Note that *D*^*x*^ is the same for all points in the domain. Elements of 𝕊^*n-*1^ are denoted by *σ* with measure d*σ*. Using Lemma 1 and assuming that the set of destinations is uniform across the domain, we can rewrite the Master equation (7).

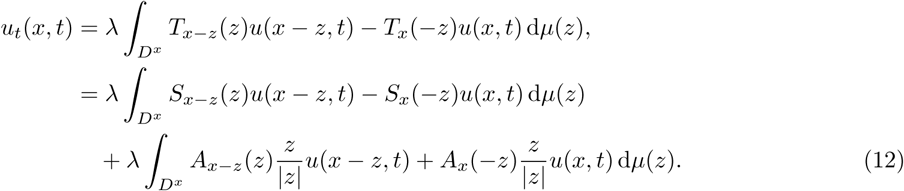

Using assumptions 1 and 2, and the fact that ‖ *σ* ‖ = 1, we can rewrite (12) as

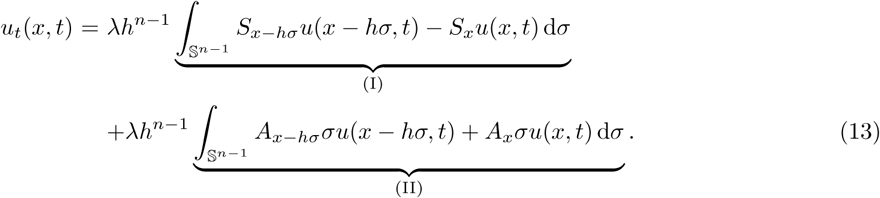

Let us consider the integrals (I) and (II) separately.

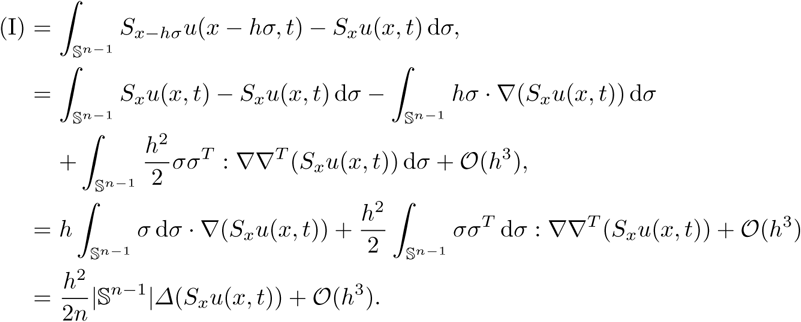

The colon notation used above denotes the contraction of two rank-two tensors as

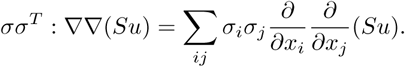

In the last step we used the fact (see for example (Hillen, 2005)) that

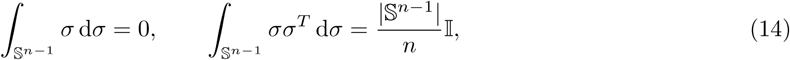

where I is the n×n identity matrix. Consider integral (II)

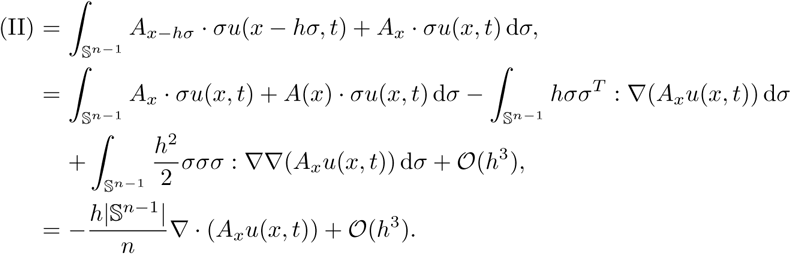

Here the colon notation denotes the contraction of two rank-three, tensors as

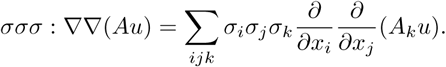

Additionally, we used the above identities (14) and the fact that

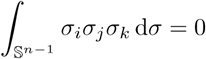

for all *i, j, k* = 1*, …, n*, as shown in (Hillen, 2005). Using the approximate expressions (dropping all 𝒪 (*h*^3^) terms) for the integrals (I) and (II) in equation (13) we obtain

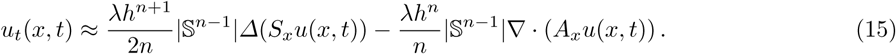

It is interesting to note that the even part of *T* enters the diffusion term, while the odd part enters the drift term.

#### 2.3 Scaling

We now study the asymptotic behaviour of equation (15). In the following *u*(. *, t*) will denote a possible non-local dependence on the population density *u*(*x, t*). Here we are interested in the limits as the lattice spacing *h* tends to zero and large time. The large time limit is obtained by letting the rate of steps per unit time *λ* tend to infinity. Different asymptotic equations are obtained, depending on the asymptotic behaviour of *λ*(*h*). We consider two cases:

1. **Drift dominated:** In this case we assume that *S*_*x*_ and *A*_*x*_ are of order 1 relative to *h* and *λ*, and that space and time scale to the same order. That is, 1*/λ ∼ h*^*n*^, which is a hyperbolic scaling. With these assumptions, we can guarantee that the following limits exist.

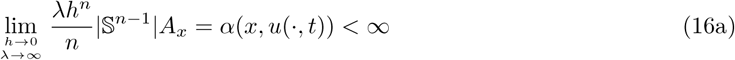

and it follows that

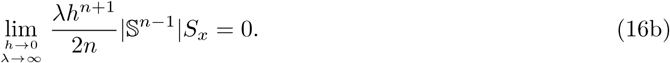 Note that in the limits (16a) and (16b), *A*_*x*_ and *S*_*x*_ respectively may depend on the cell population *u*(*·, t*). For this reason, we make the function *α* depend on *u*(*·, t*). The limit equation is then a pure drift equation

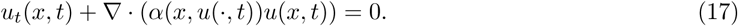
2. **Advection-Diffusion:** In this case we assume that *S*_*x*_ is of order 1 relative to *h* and *λ* and *A*_*x*_ is of order *h* and of order 1 relative to *λ*. This means, 1*/λ ∼ h*^*n*+1^, and that time is scaled with one order of *h* higher than space. This is a parabolic scaling. With these assumptions, we ensure the existence of the following limits,

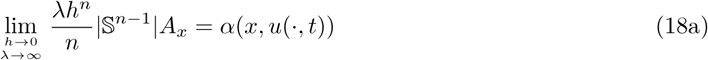

and

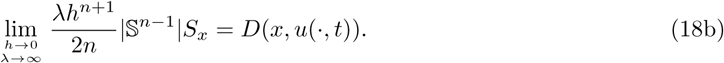 Note that in the limits (16a) and (16b), *A*_*x*_ and *S*_*x*_ respectively may depend on the cell population *u* (*·, t*). For this reason, we make the functions *D* and *α* depend on *u*(*·, t*). Then we obtain an advection-diffusion equation

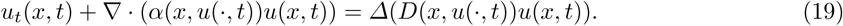 Note that the spatial diffusion constant appears inside the Laplacian: this is expected for transition rates based on local information only; for more details see (Calvo et al., 2015; Stevens and Othmer, 1997).

An in depth discussion of the different scalings together with their motivation can be found in (Hillen and Painter, 2013).

## 3 Derivation of non-local adhesion models

In this section the adhesion model (5), as proposed by Armstrong et al. (2006), will be derived using the framework developed in Section 2. In the first step we will derive an expression for the adhesive polarization vector while in subsequent steps we show how different modelling assumptions give rise to different cell-cell adhesion models. This derivation is carried out on the infinite lattice *hℤ*^*n*^. In addition we choose our time steps sufficiently small such that only a single cell moves in each infinitesimal time interval, while the remaining population effectively remains constant. This non-moving population is referred to as the *background* population in the following (see Fig. 2).

**Fig. 2:**
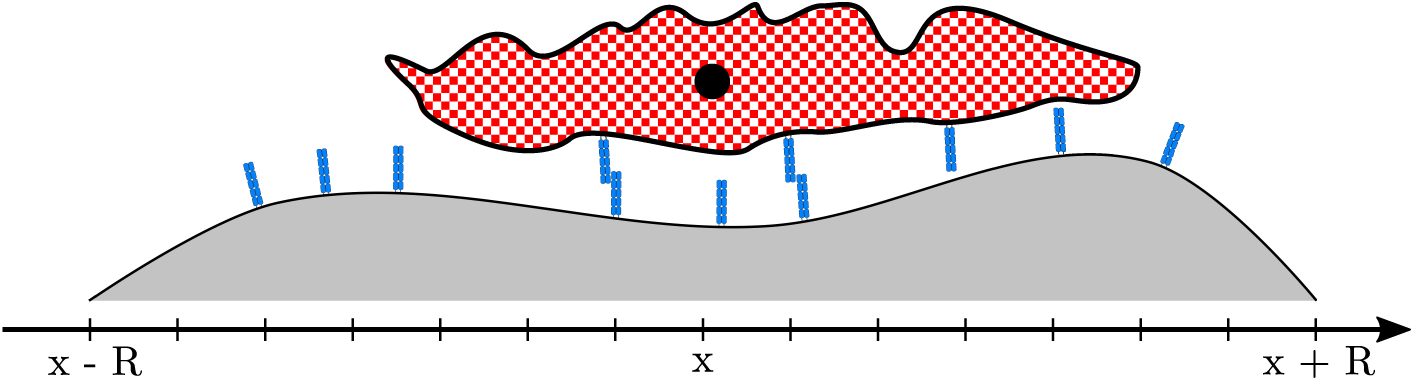
The single jumper (white-red checkerboard pattern) with its cell body (black disc) located at *x*, interacts with the background population (grey) via the adhesion molecules on the respective surfaces, represented as blue sticks.

### 3.1 Microscopic Model of adhesion molecule interactions

Consider a cell located at *x ∈ Ω*. This cell senses its environment by sending out many membrane protrusions. We now study how a single membrane protrusion interacts with the cell’ s environment in a test volume *V*_*h*_(*x* + *r*) of side length *h*, centred at *x* + *r*. We make four assumptions:

1. The probability of extending a membrane protrusion to *V*_*h*_ depends on the distance *r*. Let the proportion of the cell extended into *V*_*h*_ be denoted by *ω*(*r*). Since the protrusions are long and thin, the covered volume within the small test volume is proportional to *h*. Cell protrusions are limited in length by the cell’ s cytoskeleton, which above a certain threshold will resist further extension (Brodland and Chen, 2000). This physical limit is a natural interpretation of the sensing radius *R*. Consequently, the distribution *ω*(·) has compact support, i.e., supp *ω*(*r*) *⊂* 𝔹 ^*n*^(*R*).
2. Cell membrane protrusions are very flexible and are able to fit into very tight spaces. Nevertheless, free space is still required to establish contact. The fraction of available space in *V*_*h*_(*x* + *r*) is denoted by *f* (*x* + *r, u*(*x* + *r, t*)). This quantity is dimensionless and one choice is discussed in equation (32) later.
3. Once a cell protrusion reaches *V*_*h*_ the adhesion molecules on its surface form adhesion bonds with free adhesion molecules of the background population. If sufficiently many adhesion bonds form the membrane protrusion is stabilized and persists (Ridley et al., 2003). Otherwise, the protrusion retracts. This retraction is commonly observed as ruffles on the cell surface (Alberts, 2008). We denote the density per unit volume of formed adhesion bonds in *V*_*h*_ by *N*_*b*_(*x* + *r*). We assume that the more adhesion bonds are formed, the stronger the generated adhesion force, and the more likely it is that the protrusion persists. Later we will assume that the number of adhesion bonds in *V*_*h*_ are related to the background cell population, i.e., *N*_*b*_(*x* + *r, u* (*, t*)). Note that this dependence may be non-local as in Section 3.5.
4. The direction of the adhesion force is 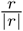

In summary the adhesion strength generated in *V*_*h*_(*x* + *r*) is determined by distance effects, free space and the number of formed adhesion bonds. Detailed functional forms for *f* (·), *ω*(·) and *N*_*b*_(·) will be discussed in subsequent sections. Let ***p***_*h*_(*x, r*) denote the adhesion strength generated in *V*_*h*_. Then, after incorporating the three effects, ***p***_*h*_(*x, r*) is given by

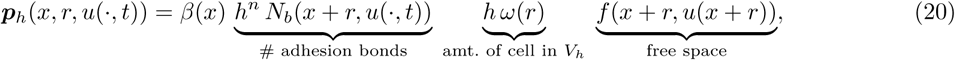

where *β*(*x*) is a factor of proportionality. The factor *β*(*x*) may include cellular or environmental properties, such as a cell’ s sensitivity to polarization.

Next we sum over all test volumes present within the sensing radius of the cell. In this step the direction of the adhesive force is important, i.e., assumption 4 above. We obtain the cell’ s polarization vector,

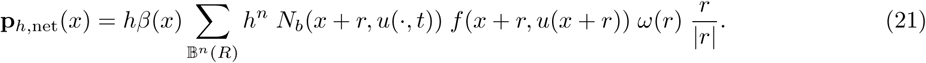

Now we can make use of our general derivation from Section 2. In detail, we let *A*_*x*_ = **p**_*h,*net_(*x*) and take the formal limit as *h →* 0 and *λ → ∞*. Note that in this case *A*_*x*_ = 𝒪 (*h*), hence limit (18a) applies and the final advection term is

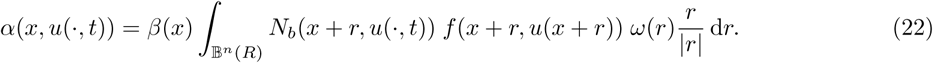

The nature of the dependence of *N*_*b*_ on the cell population *u* (*·, t*) will be discussed in the following sections. Here we do not specify *S*_*x*_ further, we only require *S*_*x*_ to satisfy the assumptions in Section 2. In particular, we will consider the case in which *S*_*x*_ does not depend on *u*(*·, t*). The complete model is then,

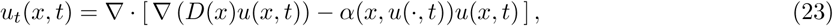

with *α*(*x*) given by (22) and *D*(*x*) by (18b).

### 3.2 The Armstrong Model

To obtain the Armstrong model (5) as a special case of (23), we assume the following choices.

1. *ω*(*r*) is the uniform distribution on the sensing region 𝔹 ^*n*^(*R*). That is,

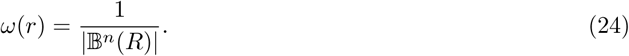
2. There is always free space and spatial constraints do not restrict the adhesion process, so *f* (*x*) *≡* 1.
3. Mass action kinetics for the reaction between the adhesion molecules of the background population and the extending cell. Specifically, let *N*_1_(*x*) and *N*_2_(*x*) denote the density per unit area of adhesion molecules of the walking cell and the background population at *x* respectively. The binding-unbinding reaction can be written

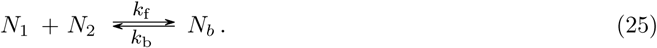 Using the law of mass action, the kinetics for this reaction are

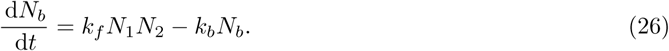 Assuming that these reactions are fast, the steady state population of *N*_*b*_ may be expressed by

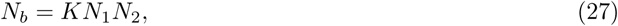 Where 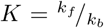. The density of adhesion molecules a given cell has on its surface depends on various factors. Here it is assumed that the density of surface adhesion molecules is a given constant. The density of adhesion molecules in the background population *N*_2_(*x*) is assumed to be directly proportional to the density of the background population at *x*. Therefore, *N*_2_(*x*) *∼ u*(*x*). With these choices for the relevant functions, equation (23) becomes

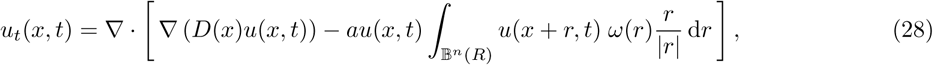

where *a* = *βKN*_1_ and *ω*(*r*) by (24). The one dimensional version of equation (28) is the adhesion model proposed by Armstrong et al. (2006).

#### 3.2.1 Different choices for ω(·)

The physical limit of cytoskeleton extensions was modelled using *ω*(·) where the compact support of *ω*(·) introduces the notion of the sensing radius. This is undoubtedly the most important contribution of *ω*(·). However, the distribution *ω*(*·*) may vary between different cell phenotypes and we briefly discuss some commonly used distributions. The simplest would be the uniform distribution, i.e.,

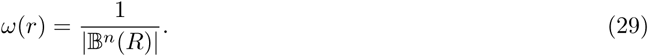

Such a uniform distribution may be too unrealistic. The resistance to extension in the cytoskeleton may build up gradually. In such a case a distribution like a triangle distribution or a Gaussian may be more appropriate. In *n*-dimensions the triangle distribution generalizes to a cone shaped distribution,

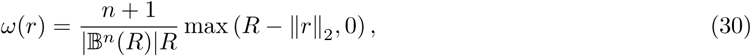

where the prefactor normalizes the distribution. For a discussion of the use of the Laplace or Gaussian distributions see (Painter et al., 2015). Mathematically, we may choose any distribution *ω*(·) which satisfies *ω*(*·*) *≥* 0 and *ω*(*·*) *∈ L*^1^(*·*). Note that these are the only required conditions on *ω*(*·*) to guarantee the existence of solution to (5) (Winkler et al., 2016).

### 3.3 Volume Filling

In this section two different mechanisms are studied by which volume filling can be included in the adhesion model.

#### 3.3.1 Destination dependent volume filling

First is the well known volume filling introduced by Painter and Hillen (2002). In this work, the transition rates are modified via a decreasing function of the occupancy of the destination site. That is, we modify the transition rates as follows:

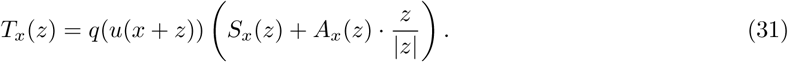

The function *q*(*·*) denotes the probability of finding space and it is chosen such that *q*(*U*_max_) = 0 and *q*(*u*) *≥* 0 *∀* 0 *≤ u ≤ U*_max_. The effect *q*(*·*) has on cell movement is that it reduces the rates of moving to areas with high occupancy. There are many possible choices of *q*(*·*). One of the most logical choices is

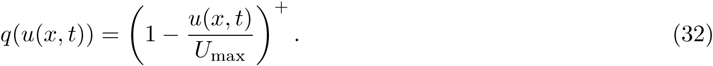

where (*·*)^+^ = max (0*,·*). For this special choice, the term modifying the diffusion term equals one (for details see Painter and Hillen (2002)). If such a volume filling is added, the adhesion model (28) becomes,

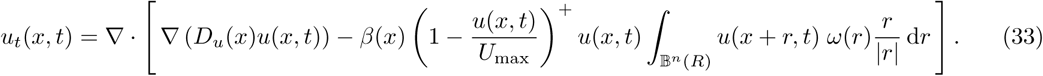

This equation describes the situation in which a cell is unable to move if it is fully surrounded by neighbouring cells.

#### 3.3.2 Adhesion molecule volume filling

A second possible form of volume filling is introduced through the free space *f* (*·*) term in the polarization of the cell (see equation (20)). The free space term in the cell’ s polarization captures the idea that high occupancy reduces the influence of that location on the cell’ s polarization: high occupancy is expected to reduce the probability of membrane protrusions that feel out that location. Secondly, a high occupancy could also correlate with a low number of free adhesion molecules. Upon letting the free space term in (20) be *f* (*x*) = *q*(*u*(*x*)), equation (23) becomes,

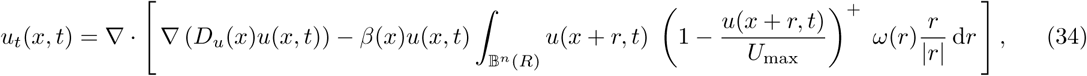

where (*·*)^+^ = max(0*,·*). Equation (34) is the same as one of the models presented by Armstrong et al. (2006), in which the *g*(*·*) function was chosen to be logistic. In this case, areas with high occupancy do not contribute to the adhesion force.

### 3.4 Bell adhesion bond kinetics

Adhesion molecules are confined to the cell surfaces and, for this reason, their binding-unbinding kinetics are different to those of a chemical species moving freely in space. This was recognized by Bell (1978) who developed detailed reaction rate constants for this situation. In this section, Bell kinetics replace the mass action kinetics (25) used in Section 3.2.

Bell kinetics are used in each of the small test volumes *V*_*h*_, in which a membrane protrusion interacts with the background population (see Section 3.1). As in equation (25) the densities of adhesion molecules on the membrane protrusion and the background population are denoted by *N*_1_ and *N*_2_, respectively. The density of formed adhesion bonds is denoted by *N*_*b*_. In contrast to mass action kinetics we distinguish now between free adhesion molecules and formed adhesion bonds. Then the densities of adhesion molecules is split,

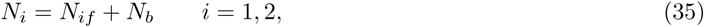

where *N*_*if*_ denotes the unit densities of free adhesion molecules. The kinetic equation governing the evolution of *N*_*b*_ is

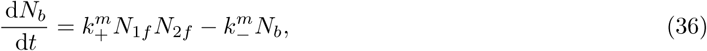

where *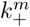* and *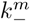* are the rate constants of bond formation and bond dissociation, respectively. For details on how the reaction rates *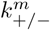* are determined see Bell (1978). To remove the dependence on *N*_*if*_ we use identity (35) to obtain,

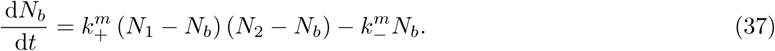

Following (Bell, 1978; Lauffenburger, 1989) it can be assumed that the number of adhesion molecules in the background population is much larger than the number of adhesion bonds formed, i.e., *N*_2_ ≫ *N*_*b*_. This is particularly true, since the number of adhesion molecules on the membrane protrusion is generally expected to be much lower than the background population, i.e., *N*_1_ *≪ N*_2_. Then equation (37) approximates to

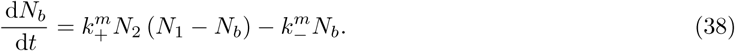

The non trivial steady state of this equation is given by

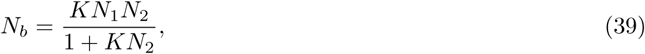

where *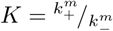* Once again it is assumed that *N*_1_ is given by a constant and that *N*_2_(*x*) *∼ u*(*x*). Then using *f* (*x*) *≡* 1 equation (28) becomes

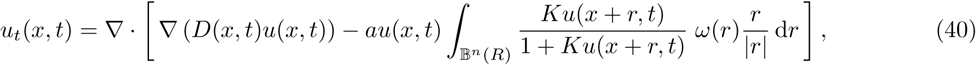

where *a* = *βN*_1_. This describes the situation in which the adhesion force saturates for large cell densities.

### 3.5 Adhesivity of the background population

In the previous sections the adhesion molecules of the background population were assumed to be directly proportional to the background population at that location, i.e., *N*_2_(*x*) *∼ u*(*x*). The implicit assumption here, however, is that adhesion molecules of background cells are concentrated at their centres. We introduce the distribution *η*(*r*) which describes the distribution of adhesion molecules across the full cell body. Then the sum total of adhesion molecules present at a location *x* is given by,

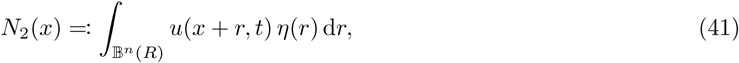

Note that *η* (*·*) has no *y*-dependence, as it is assumed to be equal for all cells. With this definition the adhesion model (23) becomes,

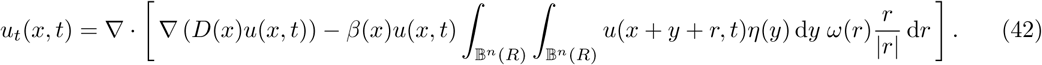

The double integral formulation of the adhesion model includes all possible adhesion molecules within a small test volume *V*_*h*_. Therefore, this formulation generalizes the adhesion model (28) by treating both the single cell jumper and the background population equal. A pictorial comparison of these two models is shown in Figure 3.

**Fig. 3:**
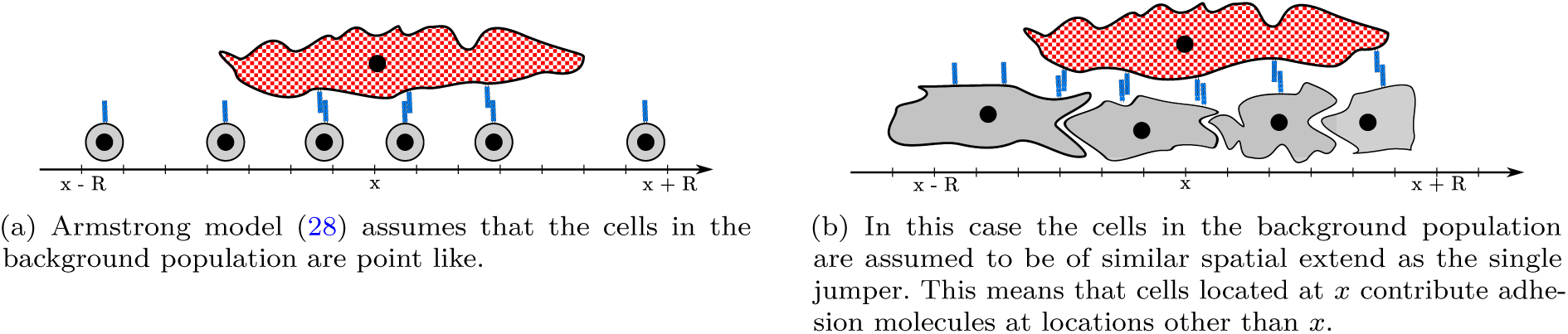
In this figure cells in the background population are shown in grey, while the single jumper is shown with a white-red checkerboard pattern. The adhesion molecules are depicted as blue sticks.

Note that, if we assume that η(r) = δ_0_(r), then we recover model (28). In summary the assumption for Armstrong’ s model (28) are that cells are point-like, have uniform densities of adhesion molecules, and that the adhesion molecules interact via mass action kinetics. At the same time, however, we retain the assumption that cells are able to sense their environment in a non-local fashion.

#### 3.5.1 Centred adhesion molecule distribution

In this section the adhesion molecule distribution *η*(*·*) is considered, in more detail. The analysis in this section will be carried out in one dimension. The main assumption of the following scaling argument is that most adhesion molecules are concentrated at the cell middle. Hence, the majority contribution to the integral in equation (41) originates around *r* = 0. Asymptotic analysis on a small parameter *∈* that controls the width of the distribution *η*(*·*) is used to simplify model (42). Introduce *∈* to control the width of the distribution *η*(*·*),

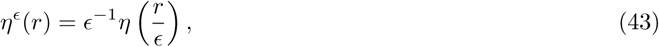

then *η*^*∈*^ integrates to unity and has compact support.

An asymptotic moment expansion can be obtained for the functional defined in (43) (Estrada and Kanwal, 1993).

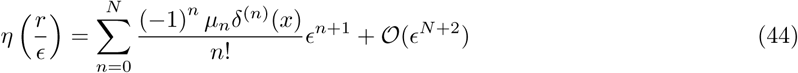

where *δ*^(*n*)^ is the *n*th derivative of the Dirac delta distribution. The moment *µ*_*n*_ is given by,

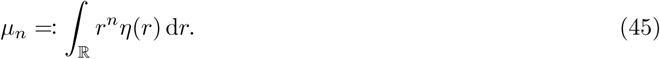

Then *η*^*E*^ becomes,

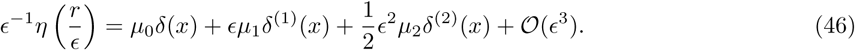

Note that *µ*_0_ = 1 as *η*(*r*) is a normalized distribution. Upon substitution of (46) (dropping all 𝒪(*∈*^3^)) into model (42) the following is obtained,

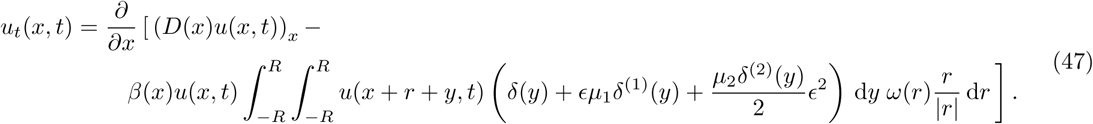

Thus resulting in

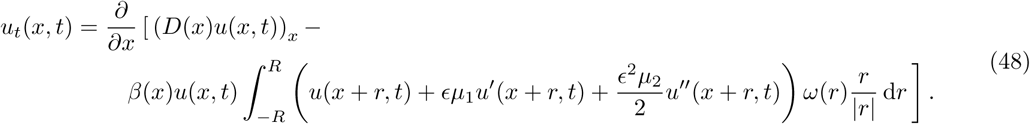

This describes the situation in which the adhesion molecule distribution on the cell surface is concentrated at the cell body. In the macroscopic equation this results in a dependence on higher order derivatives within the non-local term. An open question follows: “In what sense does this equation converge to model (28) as *∈ →* 0?”.

## 4 Derivation of the non-local chemotaxis model

Chemotaxis defines the directed movement of a biological cell in response to an external chemical gradient. A cell may either move up or down this gradient. Therefore, the directional motion component of the flux is proportional to the chemical gradient. One of the most prominent features of the Keller-Segel chemotaxis model is the formation of blow up solutions (Horstmann, 2003), although these are not biologically realistic. This has been remediated in many ways, one of which is through the introduction of a non-local chemical gradient by (Othmer and Hillen, 2002) and (Hillen et al., 2007). The non-local gradient is motivated by the observation that a cell senses the chemical gradient along its cellular surface. In *n*-dimensions the cell surface is approximated by a sphere of radius *R*, and the non-local gradient is then defined as,

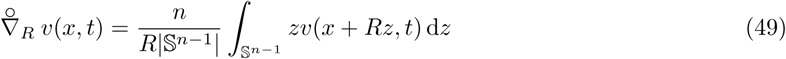

where *z* is the unit outward normal. Once again the conservation equation (1) can be used, by setting the flux *J* = *J*_*d*_ + *J*_*c*_, where *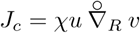*. We then obtain,

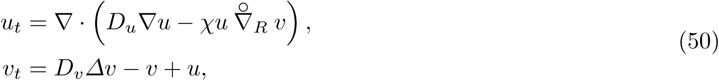

where the evolution of the chemical *v*(*x, t*) is modelled using a reaction-diffusion equation. It was shown that this model features globally existing solutions (Hillen et al., 2007).

It is well established that cells undergoing chemotaxis polarize in response to the gradient of an external chemical cue (Jilkine and Edelstein-Keshet, 2011; Weiner et al., 2000). Intracellular mechanisms then amplify, interpret and select the polarization direction (Weiner, 2002). Consequently, it is expected that the cell’ s polarization vector is proportional to the chemical gradient that the cell detects.

Following the general derivation in Section 2 we let *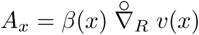*. We then obtain two different cases depending on the asymptotic behaviour of *A*_*x*_.

**Taxis dominated migration:** Suppose that *A*_*x*_ *∼ 𝒪* (1). In this case we use limit (16a) and limit (16b). to obtain,

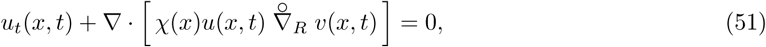

where 𝒳 (*x*) is determined by the limit (16a). Taxis dominated chemotaxis models such as this have been studied in detail in (Dolak, 2004).

**Advection-Diffusion Limit:** Suppose that *A*_*x*_ *∼ 𝒪* (*h*). In this case we use limit (18a) and limit (18b), to obtain

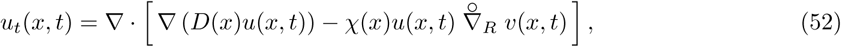

where *𝒳* (*x*) is determined by limit (18a) and *D*(*x*) is determined by limit (18b). This equation is the non-local chemotaxis model (50) from (Hillen and Painter, 2009). The key idea of this *space-jump* derivation of the non-local chemotaxis model was that the sensing radius of the cell remains constant while taking the mesh size *h* to zero.

## 5 Numerical verification of the derivation

In this section we carry out a numerical verification of the space-jump process presented in Section 2 for the adhesion model (5). For this purpose we will solve both the stochastic random walk and the partial differential equation (5) and compare the results. The stochastic simulation is implemented using the well known Gillespie SSA algorithm (Erban et al., 2007; Gillespie, 2007). The non-local partial differential equation is solved using a method of lines approach, for details see (Gerisch, 2010). Both simulations are carried out on a one dimensional finite domain, with periodic boundary conditions. For the detailed implementation of both simulations see Section 5.1 and Section 5.2.

### 5.1 Outline of the stochastic simulation

The Gillespie algorithm is originally formulated for the reactions between chemical species (Erban et al., 2007; Gillespie, 2007). However, it can be applied to spatial phenomena as well. As a foundation for our algorithm we use the stochastic diffusion process from Section 3.2 in (Erban et al., 2007). This means, the domain is discretized into uniform intervals, i.e., *Ω* = *hℤ*. The state vector *Y*_*i*_(*t*) denotes the number of cells on lattice site *i*. Then the reactions between the compartments are,

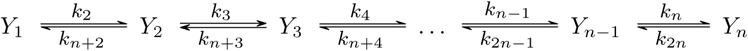

and for the periodic boundary conditions

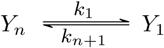

Here the reaction rates are non-local spatial functions (see definition of the rates *T* (*x, y*) in Section 2). The reaction rates for the non-local adhesion models are thus given by,

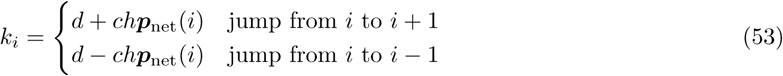

where ***p***_net_(*x*) is from equation (21). This means that ***p***_net_(*i*) at lattice location *i* is given by,

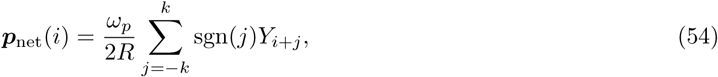

where *k* is chosen such that *R* = *kh*. Note that in this case the second factor of *h* is not required as *Y*_*i*_(*t*) already represents a population and not a density. In other words, *hN*_*b*_(*i*) ∼ *hu*(*i*) = *Y*_*i*_(*t*). The diffusion coefficient and drift coefficient are transformed into reaction rate constants *d* and *c* respectively, via limits (18b) and (18a). That is,

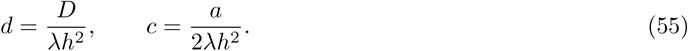

The Gillespie algorithm is implemented in C++, while the setup, data processing and plotting is written in python. The Mersenne Twister algorithm is used to generated random numbers^1^. All the numerics were carried out on an Intel Core-i7 4790K (Haswell) running Linux. The simulation parameters are given in Table 1.

**Table 1:** Parameters for Adhesion simulations

| Model Parameter | Value |
| --- | --- |
| Domain Size $L$ | 3.0 |
| Domain subdivisions per unit length | 96 |
| Diffusion coefficient $D$ | 1.0 |
| Adhesion strength coefficient $a$ | 0.003 |
| Sensing radius $R$ | 1.0 |
| Transition rate scaling $\lambda$ | 100 |
| Weighting size $\omega_p$ | 0.82 |
| Initial density $u_0$ | 4800.0 |
| Initial conditions (IC) | $u_0(1 + \kappa \sin(x))$ |
| Perturbation Size of IC $\kappa$ | 0.09 |

The implemented Gillespie SSA algorithm was verified against two test cases; a constant diffusion simulation, and a constant diffusion with constant advection simulation. The average of 64 stochastic paths generated for both test cases was compared to the solutions of the constant diffusion equation and the constant advection-diffusion equation respectively. The solutions of both test case PDEs were computed using spectral methods. For this the discrete Fourier transform methods from NumPy 1.9^2^ were used. In both cases the average of the stochastic paths agreed with the solutions of the PDEs. The verification results are not shown.

### 5.2 Outline of the numerical method for the adhesion model

For the comparison the simple Armstrong adhesion model (56) is solved on a one dimensional interval [ 0*, L*] with periodic boundary conditions. The equation is solved using a method of lines approach, for more details see Gerisch (2010).

The detailed model formulation is,

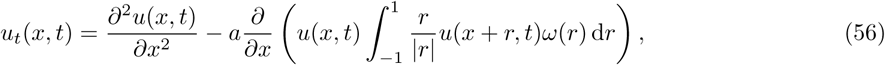

subject to

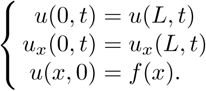

The function *ω*(*·*) is the uniform distribution over the sensing radius. That is

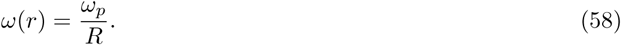

Note that the normalization factor in the continuum case ^1^*/*_*R*_ is different from the normalization factor in the stochastic simulation ^1^*/*_2*R*_ see equation (54). The simulation parameters are listed in Table 1.

### 5.3 Simulation Parameters

The parameters for both the stochastic and the continuum numerical solutions are listed in Table 1. Both the diffusion coefficient and the sensing radius were set to 1.0 to satisfy the non-dimensionalization in (Armstrong et al., 2006). The population density was not rescaled as the stochastic simulation tracked individual particles. The domain size was chosen such that only a single solution peak would form (see Fig. 4). The initial cell density was chosen such that it corresponds to a sufficiently large number of particles in the stochastic simulation. The initial density of 4800 cells per unit length, for example, corresponds to approximately 50 cells per lattice site, and a total of approximately 15000 particles. For smaller total cell numbers we observed that the results from the continuum and stochastic simulation deviated for intermediate times, while agreeing for long times. In particular, the time to reach steady state was shortened for the stochastic simulation. The adhesion strength *a* was chosen such that the constant steady state of equation (56) is unstable. Finally, the transition rate scaling *λ* was chosen and set to 100. Variations of this constant did not affect the simulation outcome.

**Fig. 4:**
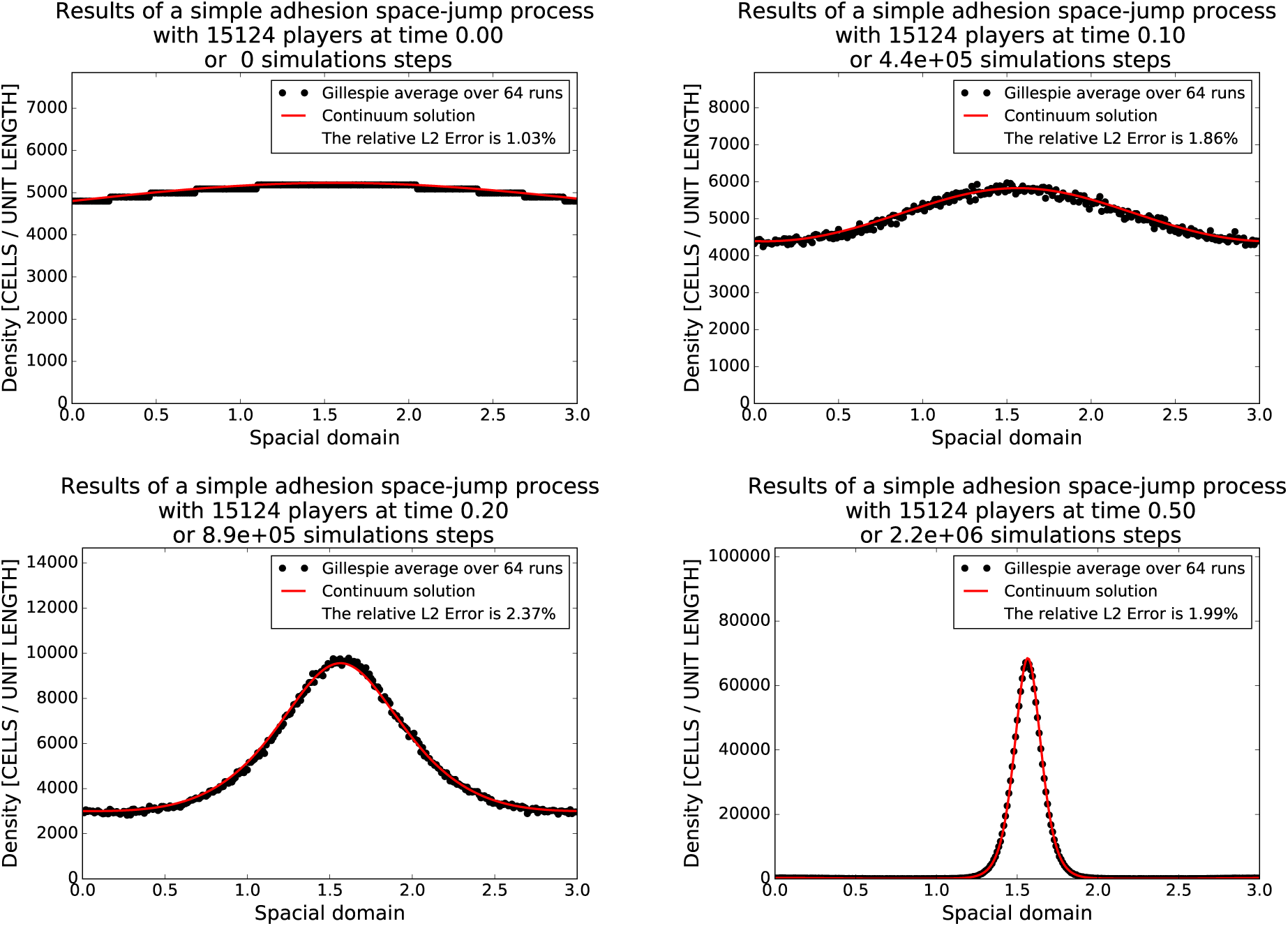
The average over 64 stochastic simulations of the state vector is shown as black circles, one for each lattice site. The line (red) represents the solution of equation (56). The first image shows the initial condition, then times 0.1, 0.2 and 0.5 are shown. Steady state is reached some time before time point 0.5. The initial condition is given by *f* (*x*) = *u*_0_ + *κu*_0_ sin(*x*). Note the stochastic paths are shifted by the procedure described in Section 5.5. For details on the implementation of the numerical simulations see the appendix. The simulation parameters are listed in Table 1.

### 5.4 Results

In Fig. 4 we compare the average density of the state vector of 64 stochastic simulations at different time points to the continuum adhesion model (56). The relative error between the average density from the stochastic simulations and the numerical differential equation solution is, at maximum, *≈*2%. Thus, under appropriate functional choices, the stochastic model converges to the Armstrong et al. (2006) model.

### 5.5 Correction of adhesion paths

Typical sets of paths generated via the Gillespie SSA with the adhesive reaction rates (53) are shown in Fig. 5a. Studying Fig. 5a it is noted that the peaks do not form at the same location between runs. This behaviour is expected, since we have a periodic domain and almost uniform initial data. This disagreement is corrected for by shifting the medians of each individual path to the location at which the PDE forms its peak. This is not a problem because the equation is solved with periodic boundary conditions. For an example of the result of such a correction see Fig. 5b. Note that the agreement between the average of stochastic simulation and the continuum solution is much better.

**Fig. 5:**
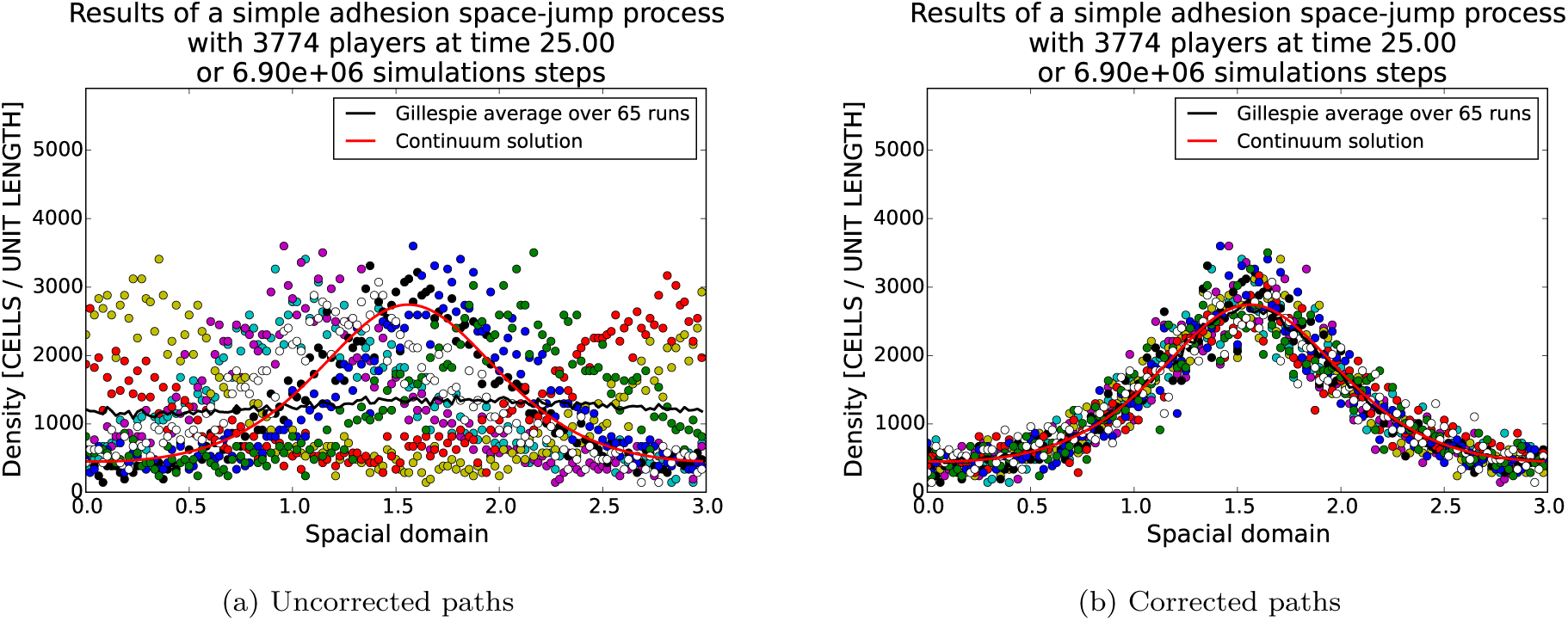
On the left a typical set of 8 simulation paths generated with the Gillespie SSA algorithm. Each separate path is indicated by a different colour (please see online version for the colour figure). The straight (red) line is the continuum prediction and the black curve the average of the 8 stochastic paths. On the right are the same paths after rotation. Rotation is done as described in the text.

## 6 Discussion

The adhesion model of (Armstrong et al., 2006) has been a step forward in the modelling of cohesive cell populations. The model has been used successfully in applications ranging from cell sorting to cancer invasion. Non-local models have been used previously to model cohesion in biological processes, see for example, the work on swarming behaviour by Mogilner and Edelstein-Keshet (1999).

Our derivation from a stochastic random walk perspective allows us to connect the macroscopic quantities of drift and diffusion to microscopic properties of cell behaviour. We analyzed adhesion binding and unbinding dynamics, the availability of free space, the extensions of protrusions and the force balances between adhesion in different directions. We found that the simplest case, in which background cells are local, free space is plenty and adhesive forces are proportional to the cell population, leads to Armstrong’ s model. However, we also show that more complicated and more realistic assumptions can be included to obtain extensions to Armstrong’ s adhesion model. It is an interesting task for future research to study the properties of volume filling, adhesion saturation and double non-locality.

The key to the presented derivation is the definition of the polarization vector. It is commonly known that cells polarize due to many different gradients (Charras and Sahai, 2014), including adhesive gradients (Nishiya et al., 2005; Ridley et al., 2003; Théry et al., 2006). Further, during embryo compaction, adhesion between neighbouring cells is established by E-cadherin dependent filopodia, which extend and attach to neighbouring cells. Subsequently, the filopodia remain under tension to bring cell membranes into close contact (White and Plachta, 2015). This observation fits closely with our definition of the adhesive polarization vector. The definition of the polarization vector allows us to keep our derivation general. This makes it straightforward to derive other taxis models through simply replacing the polarization vector. Here we derived two taxis models: the non-local adhesion model (5) and the non-local chemotaxis model (50). It is easy to envision similar derivations for other mechanisms of cell polarization.

The polarization of cells is persistent, meaning that even in the absence of an external signal the cells retain their polarization (Li et al., 2008). For instance, amoebae show a persistence time of approximately 10 min (Li et al., 2008). Li et al. (2008) conclude that the motion of amoebae is not a simple random walk, but a process by which left and right successively follow each other to avoid expensive backtracking. During the development of the *Dictyostelium* slug, periodic waves of chemo attractant cAMP are observed (Dormann and Weijer, 2001) which will result in a flipping of the gradient as it passes over the cell. Despite this, the direction of cell migration does not change (Dormann and Weijer, 2001). Such a persistence mechanism has been studied in other modelling approaches, such as velocity jump processes (Hillen, 2002). It remains, however, an open problem of how to include persistence of cell polarization within the space-jump framework.

A key assumption of our derivation is that the mean waiting time or mean residency time between jumps, is a constant with respect to the population density. An interesting direction of future research would be to weaken this assumption. In the case of cell-cell adhesion, the mean residency time would be expected to increase when strong cell adhesions are made with juxtaposed cells. Therefore, it would be expected that the mean residency time *λ* is an increasing function of cell density. A good starting point for such an investigation may be the work of (Shi et al., 2014; Zaburdaev, 2006), who both considered a coupling between the waiting time distribution and jumps.

In this work we have not been concerned with the internal cell dynamics which translate extrinsic or intrinsic cues into cell polarization. For a review discussing internal cell dynamics giving rise to cell polarization see (Jilkine and Edelstein-Keshet, 2011). A common feature of these models is a symmetry breaking process, thus distinguishing the cell front and back. It is an interesting task, to couple such a detailed model of cell polarization to our cell movement model. Such a multi-scale approach would be particularly interesting with respect to the persistence of cell polarization: for example, is it possible to obtain cell polarization by coupling two such models? A further interesting modelling question is whether such a coupling gives rise to models similar to the non-local chemotaxis or non-local adhesion model.

Mathematically, the non-local model (5) is very interesting. For the single non-local model existence and uniqueness results are available (Chaplain et al., 2011; Winkler et al., 2016) while the existence of travelling wave solutions has been demonstrated by Ou and Zhang (2013). The existence of spatially non-homogenous steady states and hence pattern formation has been numerically observed by Armstrong et al. (2006). However, an analytical treatment of the steady states of this model and the conditions under which they form remains a challenge.

The doubly non-local model (42) appears to be mathematically new. For this reason, the questions of existence, uniqueness, travelling waves, steady states are all open problems. It is further an open problem to understand the differences of such a doubly non-locality compared to the single non-locality on the form of solutions observed.

## Acknowledgements

AB was supported by NSERC, Alberta Innovates and PIMS. TH was supported by NSERC. AG thanks the Isaac Newton Institute for Mathematical Sciences for its hospitality during the programme *Coupling Geometric PDEs with Physics for Cell Morphology, Motility and Pattern Formation*; EPSRC EP/K032208/1. KJP thanks the Politecnico di Torino for a Visiting Professor position.

## Footnotes

1 Implementation from libstc++ gcc 4.9.3 <u>https://gcc.gnu.org/</u>

2 www.numpy.org

